# Absence of Coronavirus in terns in the Western Indian Ocean?

**DOI:** 10.1101/2023.06.30.547193

**Authors:** Camille Lebarbenchon, Chris Feare, Christine Larose, Solenn Boucher, Audrey Jaeger, Matthieu Le Corre

## Abstract

We investigated coronavirus circulation in three tern species, in four islands of the Western Indian Ocean (Bird, Reunion, Europa, Juan de Nova). None of the 1513 samples tested positive by RT-PCR. We discuss the implication in term of host species range, ecological drivers of virus transmission, and diagnostic tools.

---

Bats and birds are natural hosts for Coronavirus (CoV; *Coronaviridae*). The emergence of bat-related CoV in humans has stimulated research on these viruses for the past decade, but knowledge on the eco-epidemiology and evolution of bird-borne CoVs (gamma- and delta-CoV) remain limited (Wille and Holmes 2020). CoVs have been detected in wild birds on all continents, but a major sampling bias toward wild ducks has led to uneven reports of virus prevalence and diversity between bird taxa (Wille and Holmes 2020, for review). Further epidemiological studies are thus needed to better assess CoV host species range, to identify the ecological drivers of viral infection, and also their spill-over potential to poultry.

Spatial isolation could represent a major barrier to the natural introduction of infectious agents on oceanic islands, but may also generate ecological conditions favoring the local maintenance of viruses in wild bird communities inhabiting these islands. In the Western Indian Ocean, oceanic islands are major breeding sites for seabirds, with several species aggregating at very high densities. Most of these species are pelagic; migratory gulls and ducks do not breed or roost on these islands. We previously identified Brown noddies (*Anous stolidus*) and Lesser noddies (*Anous tenuirostris*) (Charadriiformes) as major Avian Influenza virus (AIV) hosts and, to a lesser extent, Sooty terns (*Onychoprion fuscatus*; Lebarbenchon et al. 2015). Major differences in the prevalence of AIV positive and seropositive birds were reported between taxa (Lebarbenchon et al. 2013, 2015), suggesting species-specific variation in AIV circulation. CoVs were also screened in > 300 cloacal swabs collected from eight seabird species, mostly from Phaethontiformes, Procellariformes and Suliformes, but none tested positive for the presence of CoV RNA (Lebarbenchon et al. 2013).

In this study, we investigated CoV circulation in three Charadriiformes species, the Brown noddy, the Lesser noddy and the Sooty tern, on four islands of the Western Indian Ocean. Bird Island, the northernmost island of the Seychelles archipelago (3°43’S, 55°12’E), is a major breeding site for terns, with approximately 400 000 pairs of Sooty terns, c. 10 000 pairs of Brown noddies and c. 19 000 pairs of Lesser noddies (Feare 1976; Feare 1979). Europa (22°23’S, 40°21’E) and Juan de Nova (17°03’S, 42°43’E), located in the Mozambique Chanel, are hosting two major Sooty tern colonies (760 000 and 2 000 000 breeding pairs, respectively; Le Corre and Jaquemet 2005). Small populations (hundreds to several thousands) of brown and lesser noddies breed and roost on Reunion Island (21°22’S, 55°34’E).

The presence of CoV RNA was looked for in samples previously collected and tested for AIV (N = 1459; (Lebarbenchon et al. 2015, 2023) as well as in other samples (N = 54), from Brown noddies and Lesser noddies on Reunion Island (Table 1). Only adult birds were included in the study. For each bird, cloacal and oropharyngeal swabs were collected and placed in a single tube, containing 1.5 ml of Brain Heart Infusion (BHI) media (Conda, Madrid, Spain) supplemented with penicillin G (1000 units/ml), streptomycin (1 mg/ml), kanamycin (0.5 mg/ml), gentamicin (0.25 mg/ml), and amphotericin B (0.025 mg/ml). Swabs were maintained at 4°C in the field, shipped to the laboratory within 48 hours, and held at -80°C until tested.

**Table 1:** Number of tested samples per each tern species, island and year.

| Species | Island | Month/Year | N tested samples |
| --- | --- | --- | --- |
| Brown noddy (*Anous stolidus*) |  |  |  |
|  | Bird | June 2012 | 33 |
|  |  | June 2013 | 90 |
|  |  | June-July 2014 | 144 |
|  |  | July 2015 | 133 |
|  | Reunion | November 2016 | 31 |
| Lesser noddy (*Anous tenuirostris*) |  |  |  |
|  | Bird | June 2012 | 32 |
|  |  | June 2013 | 90 |
|  |  | June-July 2014 | 141 |
|  |  | July 2015 | 101 |
|  | Reunion | March 2013 | 58 |
|  |  | November 2016 | 23 |
| Sooty tern (*Onychoprion fuscatus*) |  |  |  |
|  | Bird | June 2012 | 93 |
|  |  | June 2013 | 100 |
|  |  | June-July 2014 | 109 |
|  |  | July 2015 | 53 |
|  | Europa | July 2012 | 191 |
|  | Juan de Nova | June 2012 | 91 |

All samples were processed within six month after collection in the field. Briefly, samples were vortexed and centrifuged at 1500 g for 15 min. RNA was obtained with the QIAamp Viral RNA Mini Kit (QIAGEN, Valencia, CA, USA). Reverse-transcription was performed on 10 μL of RNA using the ProtoScript II Reverse Transcriptase, Random Primer 6, and RNAse inhibitor (New England BioLabs, Ipswich, MA, USA) under the following thermal conditions: 70 °C for 5 min, 25 °C for 10 min, 42 °C for 50 min and 65 °C for 20 min (Lebarbenchon et al. 2017). cDNAs were tested for the presence of the CoV RNA-dependent RNA-polymerase (RdRp) gene using a pan-CoV multi-probe real-time PCR (Muradrasoli et al. 2009) routinely used in our laboratory (Lebarbenchon et al. 2013; Joffrin et al. 2020, 2022). PCRs were performed with the QuantiNova Probe PCR Master Mix (QIAGEN, Hilden, Germany) in a CFX96 Touch^™^ Real-Time PCR Detection System (Bio-Rad, Hercules, CA, USA), with positive (Pintail CoV PBA-15 GU393339 and Bat CoV RB369 MN183188) and negative (PCR water) controls. Before RNA extraction, 10 µl of RNA of the MS2 phage was added to each sample. All samples were then tested for cDNA of the MS2 phage in order to validate the extraction and reverse-transcription steps (Ninove et al. 2011; Lebarbenchon et al. 2013).

None of the 1513 samples tested positive for the presence of CoV RdRp. Although we cannot exclude that these negative results might be associated to limited sensitivity of the PCR assay or degraded viral RNA in the original samples, this finding is consistent with our previous report that focused on other seabird taxa and was based on a lower number of tested samples (Lebarbenchon et al. 2013). Further investigations are nevertheless needed, in other seabird species and tropical islands, in order to fully decipher biological and ecological factors involved in a potential spatial restriction in CoV circulation. The development of serological tools could also provide additional information and be applied to the detection of gamma- and delta-CoV antibodies, although the immune response to CoV infection also remains to be described in seabirds (*e*.*g*. waning of CoV-specific antibodies, maternal transfer, cross-reactivity). Finally, seasonality in CoV transmission dynamics as well as differences between bird age class could also explain our negative result. Such variation in CoV shedding is suspected for ducks (Wille et al. 2015, 2017) and has been described for bats (*e*.*g*. Joffrin et al. 2022), as well as for other host-parasite systems (Altizer et al. 2006), and also suggests that further longitudinal studies in seabird populations in the tropics are needed.

We are grateful to the Savy family for their warm hospitality and support in the field work on Bird Island. We also thank Matthieu Bastien, Sophie Bureau, Muriel Dietrich, Sébastien Lefort, Bernard Rota and Julie Tourmetz, for the collection of biological material in the field. This work was funded by the “Structure Fédérative BioSécurité en milieu Tropical” and by the tutorship institutions of the UMR PIMIT.

Bird capture, handling, and collection of biological material were approved by the Center for Research on Bird Population Biology (National Museum of Natural History, Paris), the Seychelles Bureau of Standards and the Seychelles Ministry of Agriculture, Climate Change and Environment. All procedures were also evaluated and approved by the French Ministry of Education and Research (APAFIS#3719-2016012110233597v2).

